# Aging decreases osteocyte lacunar-canalicular turnover in female C57BL/6 mice

**DOI:** 10.1101/2023.12.15.571934

**Authors:** Ghazal Vahidi, Connor Boone, Fawn Hoffman, Chelsea Heveran

## Abstract

Osteocytes engage in bone resorption and mineralization surrounding their expansive lacunar-canalicular system (LCS) through LCS turnover. However, fundamental questions persist about where, when, and how often osteocytes engage in LCS turnover and how these processes change with aging. Furthermore, whether LCS turnover depends on tissue strain remains unexplored. To address these questions, we utilized confocal scanning microscopy, immunohistochemistry, and scanning electron microscopy to characterize osteocyte LCS turnover in the cortical (mid-diaphysis) and cancellous (metaphysis) femurs from young (5 mo) and early-old-age (22 mo) female C57BL/6JN mice. LCS bone mineralization was measured by the presence of perilacunar fluorochrome labels. LCS bone resorption was measured by immunohistochemical markers of bone resorption. The dynamics of LCS turnover were estimated from serial fluorochrome labeling, where each mouse was administered two labels between 2 days and 16 days before euthanasia. Osteocyte participation in mineralizing their surroundings is highly abundant in both cortical and cancellous bone of young adult mice but significantly decreases with aging. LCS bone resorption also decreases with aging. Aging has a greater impact on LCS turnover dynamics in cancellous bone than in cortical bone. Lacunae with recent LCS turnover have larger lacunae in both age groups. The impacts of aging on LCS turnover also varies with cortical region of interest and intracortical location, suggesting a dependence on tissue strain. The impact of aging on decreasing LCS turnover may have significant implications for bone quality and mechanosensation.

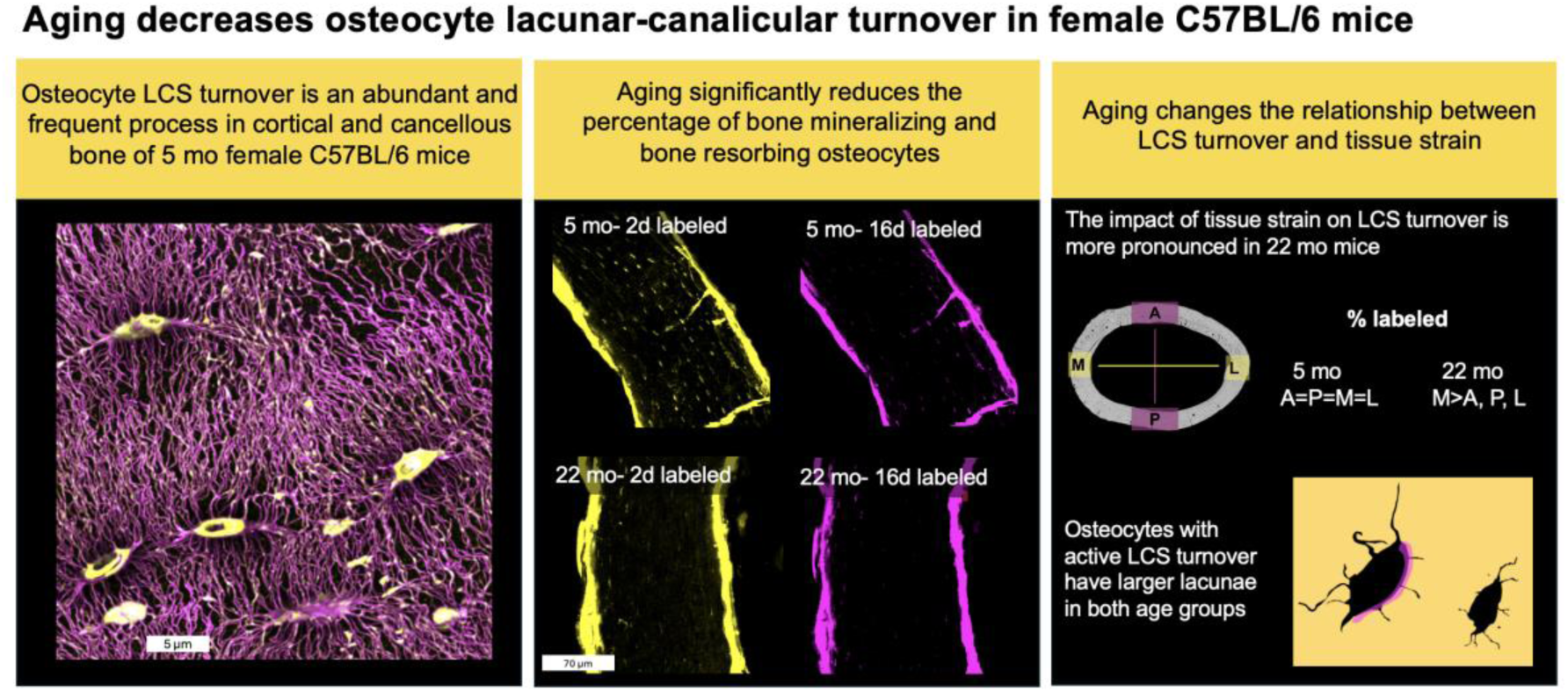
Graphical abstract. Osteocyte participation in lacunar-canalicular system (LCS) mineralization is abundant in young mice but decreases with aging in both cortical and cancellous bone. Aging also reduces LCS bone resorption. Aging alters the relationship between LCS turnover and variation in cortical tissue strain.

## 1.0 Introduction

The loss of bone fracture resistance in aging is a major public health problem^1,2^. Osteocytes, the most abundant and longest-lived bone cells^3,4^, are well-known to regulate both bone mass and bone matrix quality through the coordination of osteoblasts and osteoclasts^3–5^. The osteocyte is the topic of interest for new approaches to manage bone fragility in aging, since over time many of these cells become senescent or apoptotic^6–8^ and require greater strains to engage anabolic signaling^9–16^. Osteocytes live in a porous network within lacunae connected by canaliculi and can remove and replace (i.e., turn over) bone surrounding this network^3–5,17,18^ (**Figure 1**). There is abundant evidence that aging decreases lacunar and canalicular sizes and connectivity in both rodents and humans^6–8,14,17–22^. These geometric changes imply that bone resorption and mineralization by osteocytes alongside the lacunar-canalicular system (LCS) also shift in aging, with possible impacts to bone quality and mechanosensation^3–5,13,17^. However, many fundamental knowledge gaps persist about how osteocytes interact with their surrounding bone tissue and how these processes change in aging.

**Figure 1.**
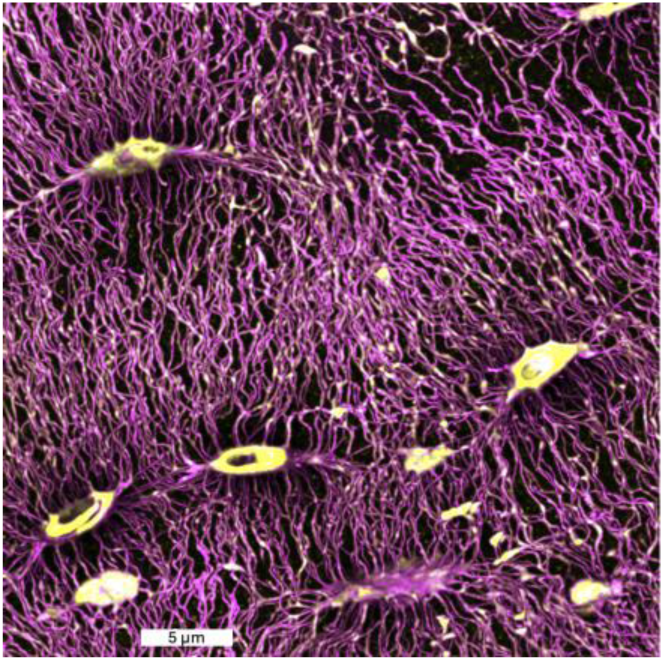
Osteocyte lacunar-canalicular system (LCS). Osteocytes live in an expansive and intricate network of lacunar holes and canalicular channels, as shown in this section of cortical femur from a 5 mo C57BL/6JN mouse. Basic fuchsin staining (magenta, *ex vivo* staining of embedded bones) shows the extensive LCS porosity of cortical bone. Calcein-stained tissue (yellow, *in vivo* fluorochrome injection 2 days before euthanasia) indicates bone mineralization.

The impacts of osteocyte LCS turnover (alternatively termed perilacunar or lacunar-canalicular remodeling) on the aging skeleton are uncertain, in part because the percentage of osteocytes that engage in bone resorption or mineralization alongside the LCS is unknown^3,4,17,18^. As witnessed in studies of rodent lactation or PTH treatment, osteocytes can expand lacunae and canaliculi through production of acids and proteases^17,18,23–33^. These pores can then recover to their original size, which implies bone re-mineralization^24,31^. When mice are injected with fluorochrome labels after weaning, abundant fluorochrome labels are observed^24,32^. However, labeled osteocyte lacunae are also seen in a number of studies where rodents were not under applied calcium pressure^30,32,34,35^, which suggests that osteocyte LCS mineralization may be a more widespread phenomena than has been previously appreciated. For example, we previously demonstrated that ∼60% of randomly-selected lacunae from the femoral midshaft cross-section were labeled with calcein administered 2 days before euthanasia in 5-month and 22-month female C57BL/6 mice^34^. In another study, ∼60% of the lacunae in the femoral midshaft cross-section of wild-type male and female C57BL/6J mice at 28-day had calcein labels administered 2 days before euthanasia^35^. Another group found that in the mid cortical cross-section of tibia, ∼55% of lacunae showed calcein labels injected 5 days before euthanasia in 2-month male wildtype littermates of MMP13 knockout mice with a mixed C57BL/6 genetic background^30^. These studies differ in age, label dosage, time of injection, region of evaluation, and mouse genetic background. Furthermore, while cortical and cancellous bone differ in their metabolic activities^36–40^, whether osteocyte LCS turnover activity varies between these compartments is unknown. Defining the percentage of osteocytes participating in LCS bone mineralization and resorption in young adult and aged mice is needed to move forward with evaluating the premise that changes to LCS turnover could impact bone quality and mechanosensation in aging.

The dynamics of LCS turnover are also essential to defining the potential impact of the osteocyte on its surrounding bone. These dynamics have been challenging to study, since LCS bone mineralization and resorption require different bone preparation and analyses. The percentage of osteocytes participating in LCS bone mineralization can be monitored by fluorochrome labeling^17,18,29–35^ (**Figure 1**). The percentage of osteocytes resorbing bone is instead measured through immunohistochemical markers of matrix metalloproteinases and other targets^19,30,35,41,42^ **(Figure 3**). However, there has been a need for an approach to estimate the dynamics of bone turnover from the same bone sections. Serial fluorochrome label injections at short intervals before euthanasia can help to address this gap in knowledge (**Figure 2**). It is not possible to assess whether an individual mouse had labeled bone that was later removed. However, the average percentage of lacunae showing labels administered at specific time points (e.g., 2 through 16 days) can allow estimation of how long labels persist following deposition. Furthermore, when serial labels are delivered to the same mice, double labels can provide an indication of how long LCS mineralization can occur (**Figure 2**). Double labels have thus far only been quantified in lactation studies as evidence of bone infilling following the removal of calcium pressure with weaning^24^. To interpret these LCS turnover dynamics, it is also necessary to assess whether common fluorochrome labels (i.e., calcein and alizarin) show similar retention around osteocyte lacunae.

**Figure 2.**
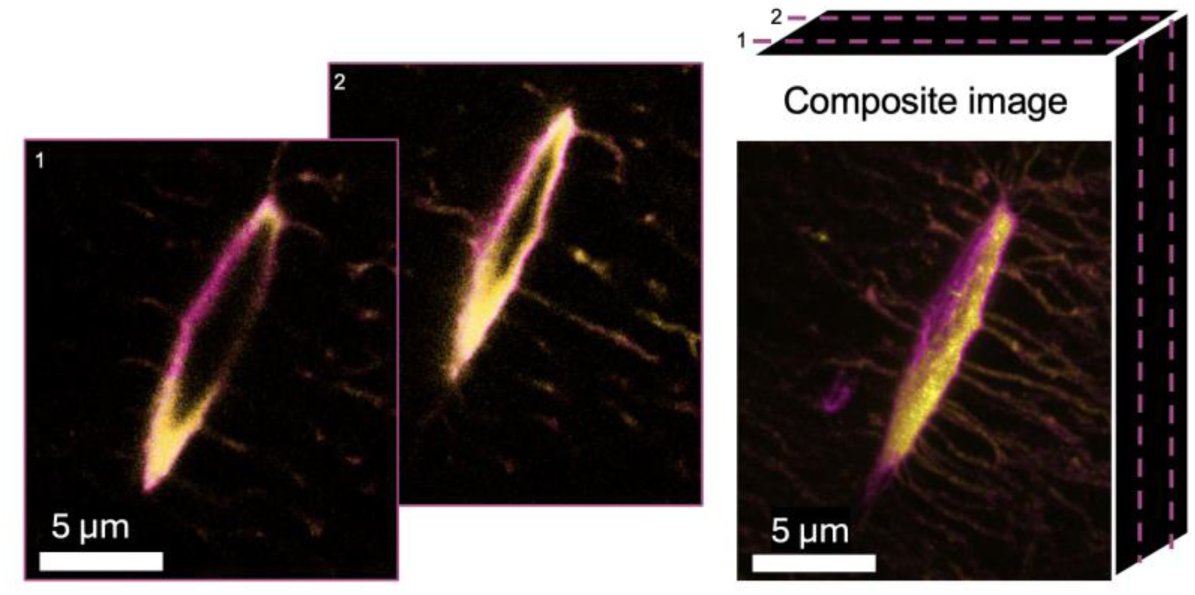
Osteocyte lacunae can show double labels when administered at short timepoints before euthanasia. *In vivo* serial fluorochrome labeling (calcein in yellow, 2 days before euthanasia; alizarin in magenta, 8 days before euthanasia) in a female 5 mo C57BL/6JN mouse reveals double-labeled lacunae.

**Figure 3.**
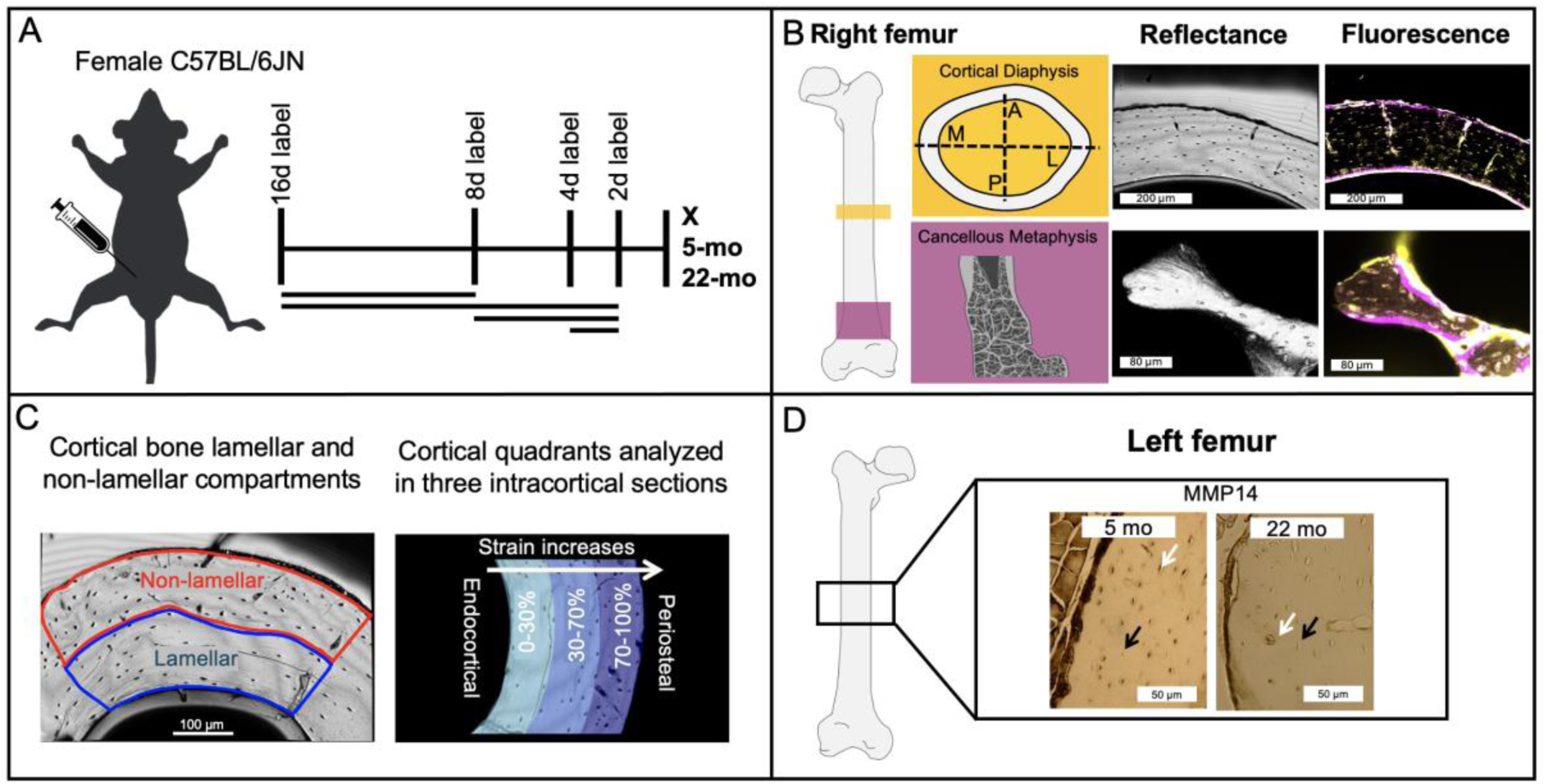
Schematic of study experimental procedures. A) Each group of mice received intraperitoneal injections of fluorochrome labels at two specific dates (16d, 8d, 4d, or 2d) before euthanasia at either 5 months or 22 months of age. B) The femur diaphysis (A/P/M/L regions) and distal metaphysis were imaged with a confocal microscope in fluorescence and reflectance modes to evaluate labeled and unlabeled lacunae on the bone surface. Representative images show 2d calcein labels (yellow) and 8d alizarin labels in (magenta) for 5 mo mice. C) Cortical bone was divided into lamellar and non-lamellar compartments for further analyses. Cortical bone was also divided into three intracortical sections for analysis based on variation in tissue strain. D) MMP14+ lacunae were counted to quantify perilacunar bone resorption for cortical bone. White arrows show MMP14+ lacunae and black arrows show unlabeled lacunae.

Another question is whether osteocyte LCS turnover is mechanosensitive and if the dependence of this process on tissue strain changes with aging. Osteocytes are mechanosensitive cells and their signaling activity depends on tissue strain^3,4,9,43,44^. Further, osteocytes are less mechanosensitive in aging and require greater strains to engage anabolic signaling^9–16^. LCS turnover has the potential to influence osteocyte mechanosensation through altering the shape of the porous LCS network as well as the flow characteristics within it^45–52^. Furthermore, as shown in our recent work using atomic force microscopy, recent LCS mineralization increases the compliance of bone within several hundred nanometers of LCS walls^34^, which would likely contribute towards strain amplification^52^. However, whether LCS turnover influences, or is influenced by, strains experienced by the osteocyte is not yet understood. Since long bones experience tissue strains that vary in direction and magnitude^53–55^, determining how LCS turnover varies throughout femoral cross-sections can allow answering important first questions about whether this process is associated with tissue strain in young adult and aged mice.

The purpose of this study is to test the hypothesis that aging decreases the percentage of osteocytes engaged in LCS bone mineralization and resorption and alters the dynamics of these processes. This hypothesis was tested by serial fluorochrome labeling and immunohistochemistry studies of cortical and cancellous compartments of the femur in 5-month and 22-month female C57BL/6JN mice. We further hypothesized that the LCS turnover activity depends on variation in tissue strain and that this relationship changes in aging, which was tested by comparing LCS bone mineralization dynamics between femur regions of interest and with distance from the endocortical to periosteal surfaces.

## 2.0 Materials and Methods

### 2.1 Animals

All animal procedures were approved by Montana State University’s Institutional Animal Care and Use Committee. Young adult 5-month-old (5 mo, n = 20) and early old age 22-month-old (22 mo, n=16) female C57BL/6JN mice from the National Institutes of Aging colony were utilized in this study. Mice had a minimum of two weeks to acclimatize to the MSU animal facility before the beginning of the label studies. Mice had *ad libitum* access to water and standard chow. Each mouse received two intraperitoneal injections of fluorochrome labels, calcein (20 mg/kg, i.p.) and alizarin (30 mg/kg, i.p.) at two specific dates. These dates included 16d, 8d, 4d, or 2d before the euthanasia. The injections were administered in a manner where each mouse received one injection of alizarin and one injection of calcein, but the specific timing and sequence of the injections varied for each group (**Figure 3A**). To ensure that label identity was not confounded with the specific time points, some mice received the calcein injection first and the alizarin injection second. Other mice received label injections in the opposite order. No significant effects of label type (i.e., alizarin or calcein) were observed on the percentage of labeled lacunae (**Figure S1**). Therefore, we aggregated the label identity data, pooling alizarin and calcein data for each label date (n = 4-16 age/label date group). Mice were euthanized at 5 or 22 months of age via isoflurane inhalation followed by cervical dislocation.

### 2.2 Sample preparation

Right femurs were harvested, cut transversely in half at the femoral midshaft using a low-speed diamond saw (Isomet, Buehler), and then the proximal and distal fragments were embedded in non-infiltrating epoxy (Epoxicure 2, Buehler) without ethanol dehydration. The proximal side of the mid-shaft cross-section was used for cortical bone studies. The embedded distal halves were cut through the sagittal plane to expose femoral metaphysis for cancellous bone studies. All samples were polished to achieve a mirror-like finish, using wet silicon carbide papers (600 and 1000 grits, Buehler) followed by Rayon fine cloths and alumina pastes (9, 5, 3, 1, 0.5, 0.3, and 0.05 μm, Ted Pella, Inc.).

Left femurs were harvested and immediately fixed with 10% neutral-buffered formalin, decalcified with EDTA disodium salt dihydrate, dehydrated in a graded ethanol series, embedded in paraffin, transversely cut at femoral midshaft, and each half was serially sliced into 5-micron-thick horizontal cortical diaphysis sections for immunohistochemistry analyses.

### 2.4 Analysis of bone-mineralizing osteocytes

An inverted confocal laser scanning microscope (CLSM-Leica Stellaris DMI8, Wetzlar, Germany) was used to identify labeled and unlabeled osteocyte lacunae using a 20X lens in air. Calcein labels were visualized using an excitation wavelength of 488 nm and emission wavelength of 500–540 nm. Alizarin labels were visualized using an excitation wavelength of 561 nm and emission wavelength of 600–645 nm. The reflectance mode was used to image the bone surface, allowing measurement of the total number of lacunae. Fluorescence and reflective images from the same site were used to calculate the percentage of fluorochrome-labeled lacunae on the bone surface (**Figure 3B**). All images were collected in z-stacks (∼30 μm thickness, 0.6 μm between slices) to confirm whether the surface-visible lacunae were labeled or not in 3D space. For each channel (alizarin and calcein) in every image, we calculated the mean and standard deviation of the grayscale intensity using ImageJ. Then, we determined a minimum intensity value by adding 1.5 times the standard deviation to the mean grayscale intensity. Using Imaris 9.3, we set the minimum threshold of fluorescent intensity for each channel in each image to this calculated value.

The percentage of bone-mineralizing osteocytes (i.e., labeled lacunae) was measured for cortical bone within anterior (A), posterior (P), medial (M) and lateral (L) regions of interest (ROIs) at the femoral midshaft (**Figure 3B**). The percentage of bone-mineralizing osteocytes was quantified for lamellar and non-lamellar compartments, as visually identified from reflectance images, for each ROI (**Figure 3C**). Cortical ROIs were also divided into three sections with respect to intracortical position with relation to the endocortical surface: 0-29%, 30-70%, and 71-100% of the cortical thickness (**Figure 3C**). For analyses of cancellous bone, multiple trabecular regions were selected from the metaphysis of each mouse (**Figure 3B**). Custom Matlab code was employed for these analyses.

During data collection, the laser for the Leica Stellaris DMI8 was updated from a diode laser to a white light laser while the detectors remained unchanged. Emission and excitation ranges were kept similar, but adjustments were made to the new laser’s settings such as gain and power to ensure a uniform image production. All images, both pre– and post-update, were normalized to their respective mean and variable intensity, as previously described. Our analysis revealed no discernible impact of the laser change (included as a blocking factor in all the statistical models) on any outcomes.

### 2.5 Analysis of osteocyte matrix metalloproteinase expression by immunohistochemistry

Paraffin-embedded left distal femurs were used for immunohistochemistry (IHC) following previously published protocols^56,57^. Slides were dewaxed and rehydrated (ethanol and distilled water series). Subsequently, the slides were incubated in Innovex Uni-Trieve retrieval solution (329ANK, Innovex Animal IHC kit) for 30 min in a 65 °C water bath. Slides were blocked with the Innovex kit’s Fc-block and Background Buster, each for 45 minutes in the room temperature. Next, samples were incubated with the primary antibodies (1:100 anti-MMP14; ab38971 both diluted in PBS) for one hour at room temperature and subsequently with secondary antibodies (Linking Ab, 329ANK) and peroxidase (HRP) enzyme for 10 minutes each at room temperature. Following this, the slides were treated with DAB working solution at room temperature for 5 minutes, washed with PBS, and mounted with Innovex Advantage permanent mounting media. Negative controls were conducted by replacing the primary antibody with rabbit IgG at the same concentration. Images were captured using a Nikon E-50i microscope (Nikon, Melville, NY, SA) with 4x (full cortical cross-section) and 20x (each cortical ROI) objectives. The mean percentage of MMP14-positive osteocyte lacunae for each cortical ROI was quantified using ImageJ (**Figure 3D**).

### 2.6 Analysis of lacunar geometry via scanning electron microscopy

A subset of samples (n=6 per age) was coated with a thin layer of carbon for lacunar geometrical analyses via backscattered scanning electron imaging (Zeiss Supra 55VP field emission SEM, 20 kV, 60 μm aperture size, 400× magnification, and 9.1 mm working distance). Samples were mounted in a custom holder that ensures flat surfaces at the same height^58^. Images were collected for the anterior ROI for lamellar bone. A custom Matlab code was used to calculate the following parameters: lacunar porosity (%, pores smaller than 150 pixels^2^ were considered lacunae), lacunar number density (#/ mm^2^), lacunar area (μm^2^), lacunar major and minor axes (μm), and lacunar circularity (i.e., ratio of minor to major axis of the fitted ellipse, where a value of 1 indicates a perfect circle). We assessed differences in lacunar geometry for labeled and unlabeled lacunae for young (n=4) and old (n=3) samples that were labeled 8 days and 2 days before euthanasia. We overlapped the SEM images with CLSM maps of labeled and unlabeled lacunae to test whether lacunar area differs between bone-mineralizing and non-bone-mineralizing osteocytes.

### 2.7 Statistical analyses

Mixed model ANOVA, with mouse as the random effect, tested whether percentage of bone-mineralizing osteocytes depended on the fixed effects of age, tissue strain (i.e., A/P/M/L ROIs), label date, or interactions of these variables for cortical bone. Additional mixed model ANOVAs were utilized for lamellar and non-lamellar cortical compartments. For the comparisons of intracortical distances (0-30%, 30-70%, and 70-100%), the two age groups were separated and for each age, mixed model ANOVA with mouse random effect was used to test if the percentage of bone-mineralizing osteocytes (for 2d and 16d labels, separately) depended on the fixed effects of ROI, intracortical distance, or their interactions. For cancellous bone, two-way ANOVA was employed to test if age, label date, and the interaction of these factors affect percentage of bone-mineralizing osteocytes. Since the confocal laser was changed mid-study, this was included as a blocking factor in these models. Because the laser change was not a significant effect for any measure, this blocking factor was removed, and models were rerun. Mixed model ANOVA with mouse as a random effect was used to test if the percentage of MMP14+ lacunae depends on age, ROI, or their interactions. For all models, model residuals were checked for satisfaction of assumptions of normality and homoscedasticity. Dependent variables were log-transformed if necessary to satisfy these assumptions. Significance was set a priori to p < 0.05. Significant interactions between factors were followed up with Tukey post hoc tests. All analyses were performed with Minitab (20).

## 3.0 Results

### 3.1 Aging decreases the number of osteocytes participating in LCS turnover

Osteocyte participation in mineralizing their immediate surrounding was highly abundant in the cortical and cancellous femur of young adult C57BL/6 mice, with more than 80% of lacunae showing 2d labels. Compared to 5 mo mice, 22 mo mice had a large decrease in the percentage of recently bone-mineralizing osteocytes (i.e., 2d labeled lacunae) in both cortical (22 mo vs 5 mo: –58%, p < 0.001, **Figure 4A & C**) and cancellous bone (22 mo vs 5 mo: –32%, p < 0.001, **Figure 4B & D**). The percentage of double-labeled lacunae (i.e., positive for both 2d and 16d labels) was also abundant in 5 mo femurs, with more than 45% of the lacunae in cortical bone and 60% of lacunae in cancellous bone having double-labels. The percentage of double-labeled lacunae decreased with aging. There were 45% (p = 0.05) and 85% (p < 0.001) fewer double-labeled lacunae in cortical and cancellous bone of 22 mo mice compared to 5 mo mice, respectively (**Figure 2**, **Figure S2**). The percentage of MMP14+ lacunae (i.e., positive for a marker of bone resorption) was also highly abundant in femoral cortical bone of 5 mo mice. For these young adult mice, more than 75% of cortical lacunae were positive for MMP14 (**Figure 4E & F**). With increased age, there was a significant decrease in the percentage of MMP14+ lacunae (22 mo vs 5 mo: –10%, p < 0.001).

**Figure 4.**
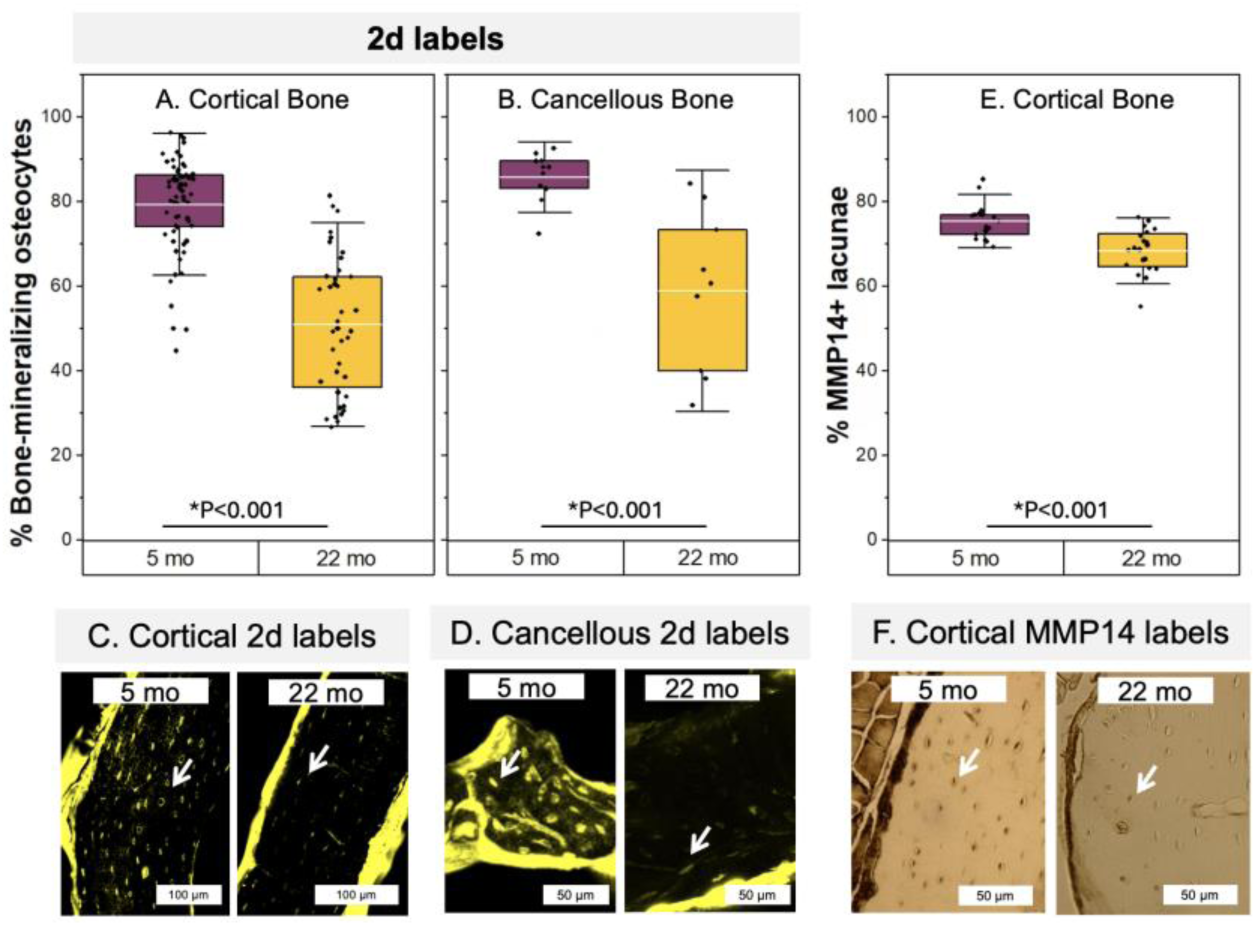
The effect of aging on cortical and cancellous osteocyte bone mineralization and resorption. Bone-mineralizing osteocytes were less abundant with aging in both A) cortical and B) cancellous bone. Only 2d labels are shown here; full time series data are provided in Figure 5. C & D) Representative fluorescence images of calcein labels for 5 mo and 22 mo mice shown. E & F) Aging also decreased the percentage of MMP14+ lacunae in cortical bone. White arrows show 2d labeled or MMP14+ lacunae. All data are reported as percentages (labeled lacunae/all lacunae). Boxplots represent mean value (cross), interquartile range (box), minimum/maximum (whiskers), and symbols representing all data points. All p-values correspond with results of the omnibus ANOVA test. * indicates a significant effect of age.

### 3.2 Aging alters LCS turnover dynamics more for cancellous than for cortical bone

The percentage of labeled lacunae decreased for injection dates further from euthanasia. For cancellous bone from 5 mo mice, the percentage of 16d labeled lacunae was 29% (p < 0.001) lower compared to 2d labeled lacunae (**Figure 5B & D**). By contrast, at 22 mo, the percentage of 16d labeled lacunae was 81% (p < 0.001) lower compared to 2d labeled lacunae (**Figure 5F & H**), suggesting that the rate of label disappearance is higher with increased age in cancellous bone. In cortical bone of 5 mo mice, there were 44% (p < 0.001) fewer lacunae labeled at 16d compared to 2d (**Figure 5A & C**). For 22 mo mice, there were 61% (p < 0.001) fewer 16d labeled lacunae compared to 2d labeled lacunae (**Figure 5E & G**), implying that in cortical bone, LCS turnover undergoes a more modest change with aging.

**Figure 5.**
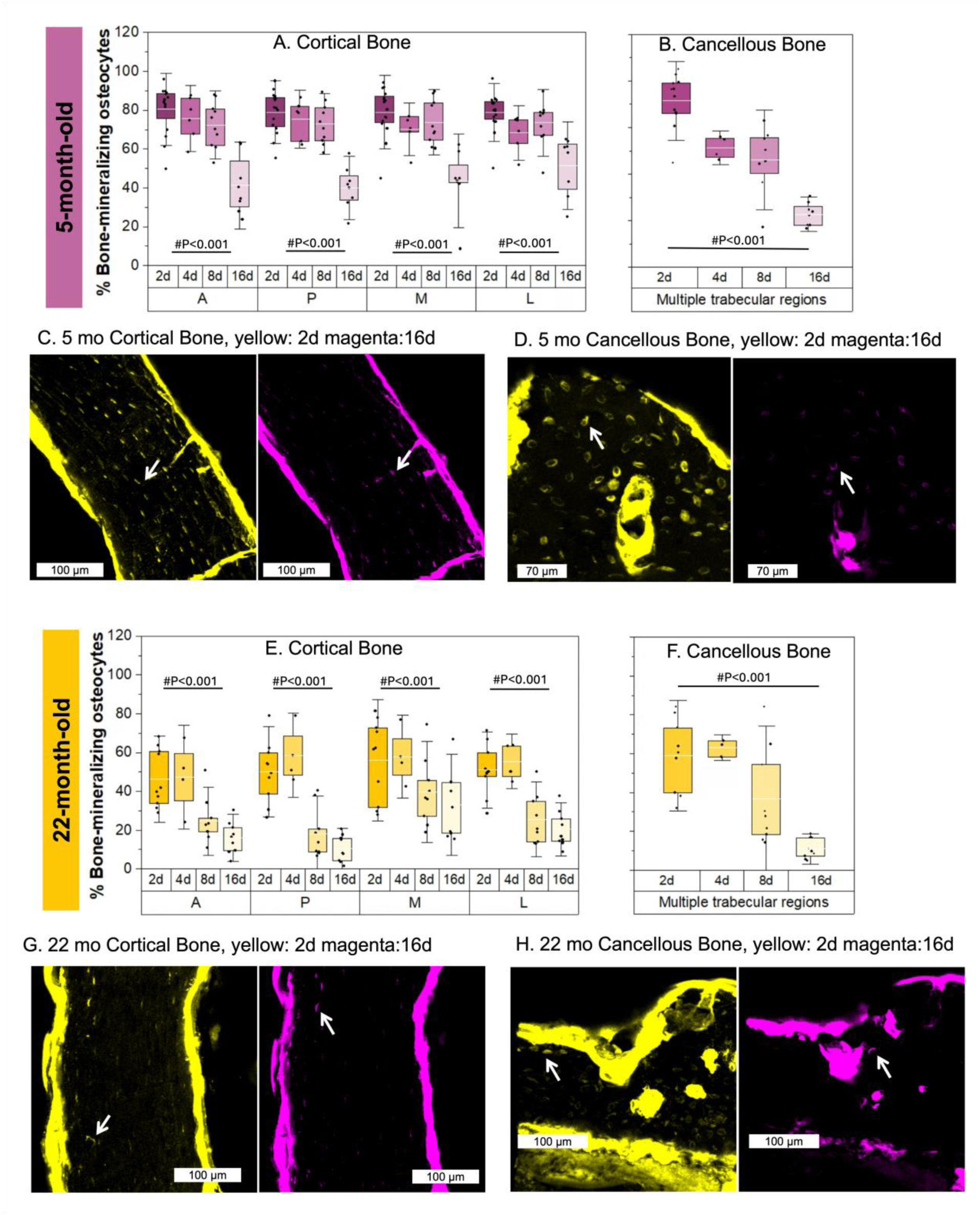
The effect of aging on the dynamics of osteocyte bone turnover. Compared to 2d labels, labels administered 16 days before euthanasia were less abundant in A & C) cortical and B & D) cancellous bone of 5 mo mice. In 22 mo mice, there were also fewer lacunae with 16d labels compared to 2d labels in both E & G) cortical and F & H) cancellous tissues. All data are reported as percentages (labeled lacunae/all lacunae). Representative fluorescence images of 2d labels (calcein label is shown in yellow) and 16d labels (alizarin label is shown in magenta) for young and old samples shown in both cortical and cancellous tissues. White arrows show 2d labeled lacunae. Boxplots represent mean value (cross), interquartile range (box), minimum/maximum (whiskers), and symbols representing all data points. All p-values correspond with results of the omnibus ANOVA test. # indicates a significant effect of injection date.

### 3.4 The impact of aging on LCS turnover is similar between lamellar and non-lamellar compartments of cortical bone

Since osteocyte lacunar geometry depends on the bone structural organization (e.g., lamellar vs non-lamellar bone) ^59–61^, we further divided the cortical bone into lamellar and non-lamellar compartments. We observed a similar age-induced decline in the percentage of bone-mineralizing osteocytes in both lamellar (22 mo vs 5 mo: –49%, p < 0.001) and non-lamellar (22 mo vs 5 mo: –46%, p < 0.001) compartments (**Figure 6A-B** for medial ROI, **Figure S3** for other ROIs). Similarly, in both age groups, 16d labels were significantly less abundant compared to 2d labels in both lamellar (5 mo 16d vs 2d: –40%, 22 mo 16d vs 2d: –62%, all p < 0.001) and non-lamellar compartments (5 mo 16d vs 2d: –45%, 22 mo 16d vs 2d: –60%, all p < 0.001) of cortical bone (**Figure 6A-B, Figure S3**).

**Figure 6.**
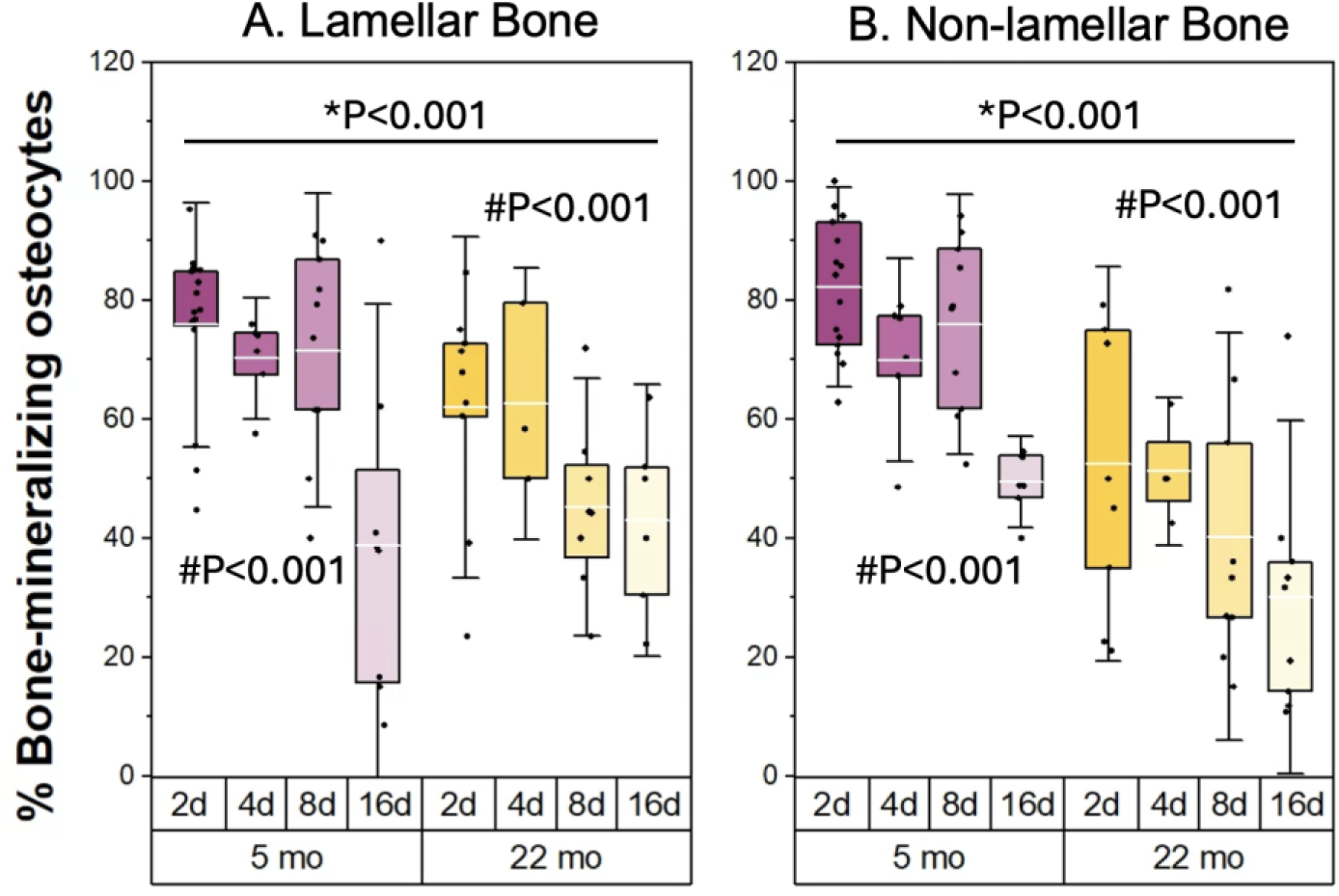
The effect of aging is similar in reducing LCS bone mineralization for both lamellar and non-lamellar bone. For both A) lamellar and B) non-lamellar compartments of cortical bone, aging reduced the percentage of bone-mineralizing osteocytes. For both types of tissues, 16d labeled lacunae were significantly less abundant compared to 2d labeled lacunae regardless of the age group. Data are shown here in the medial ROI, where every mouse had both lamellar and non-lamellar compartments. Results for all ROIs are in Figure S3. All data are reported as percentages (labeled lacunae/all lacunae). Boxplots represent mean value (cross), interquartile range (box), minimum/maximum (whiskers), and symbols representing all data points. All p-values correspond with results of the omnibus ANOVA test. * indicates a significant effect of age. # indicates a significant effect of injection date.

### 3.5 The effect of cortical tissue strain on LCS turnover depends on age

Because cortical bone experiences a range of strains across intracortical femur and the osteocyte is a mechanosensitive cell ^53–55^, we compared LCS bone mineralization and resorption for anterior (A), posterior (P), medial (M), and lateral (L) ROIs. The impact of ROI on LCS turnover depended on age (age and ROI interaction p < 0.001) (**Figure 7A**). In 5 mo mice, ROI had no significant impact on bone-mineralizing osteocytes (p = 0.815). In 22 mo mice, osteocytes in the medial ROI (close to femur neutral axis) had higher participation in mineralizing their surrounding compared to those in the other three ROIs (e.g., M vs A: +44% p<0.001). There were no differences in the percentage of labels among anterior, posterior, and lateral ROIs. Notably, the decay of 16-day labels exhibited a slower pace in the medial ROI in comparison to the others. ROI did not impact the percentage of MMP14+ lacunae in 5 mo or 22 mo mice (p = 0.7).

**Figure 7.**
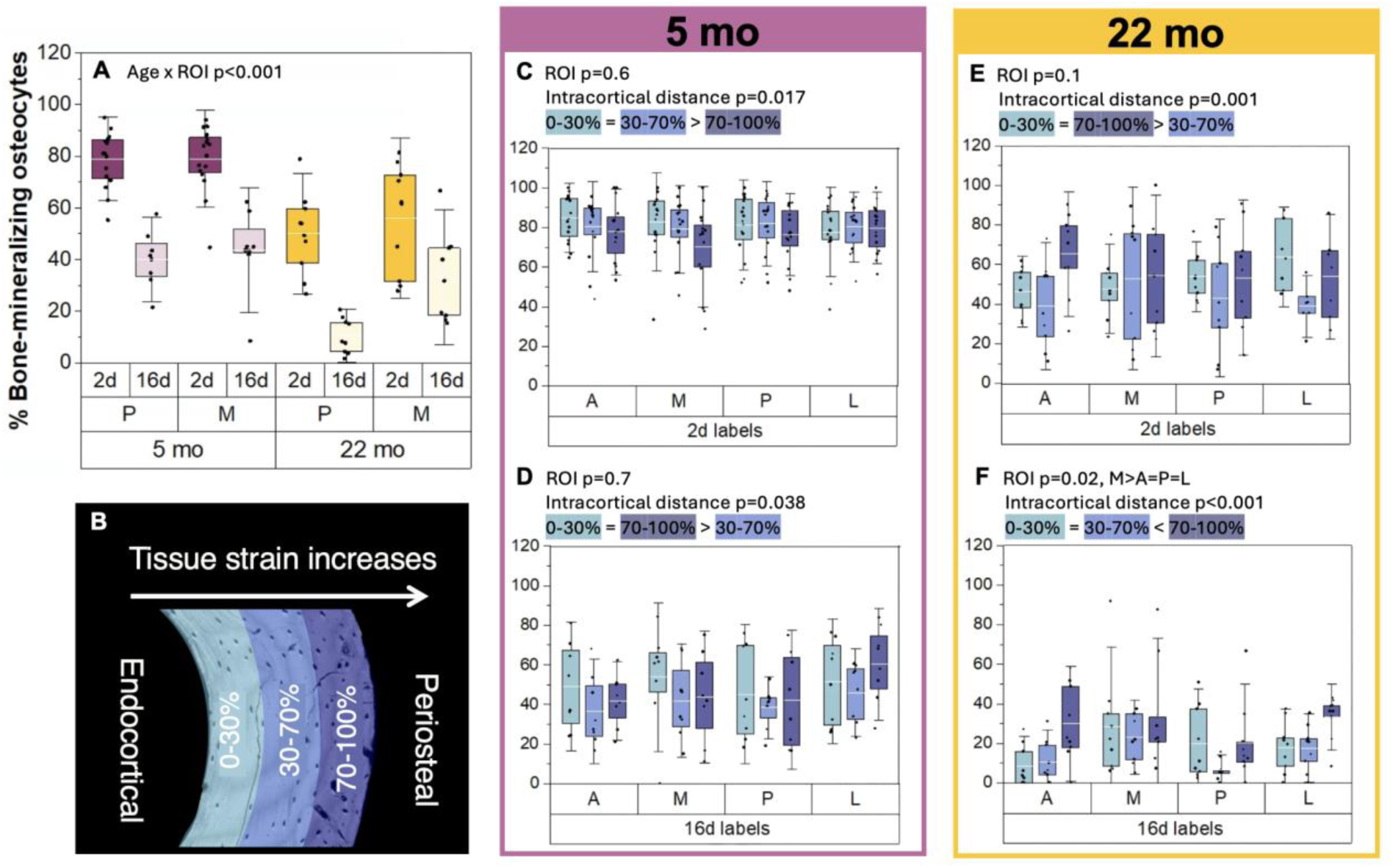
Associations between tissue strain and osteocyte participation in LCS bone mineralization. A) In 5 mo mice, cortical ROI did not impact the percentage of bone-mineralizing osteocytes, however in 22 mo mice, lacunae in the medial ROI had the highest percentage of labeling compared to all ROIs. B) Each cortical ROI was divided into three distance sections from the endocortical surface (0-30%, 30-70%, and 70-100% of cortical thickness) for further investigation of whether osteocyte bone mineralization is associated with differences in tissue strain. C) In 5 mo mice, the percentage of 2d labeled lacunae was lowest in the region closest to periosteal surface. D) For 5 mo mice, the percentage of 16d labeled lacunae was lowest in the middle intracortical region. E-F) In 22 mo mice, the percentage labeling differences between intracortical sections were larger. E) Middle intracortical section in 22 mo mice had fewer 2d labeled lacunae compared to other sections and F) the percentage of 16d labeled lacunae was highest in the region closest to periosteal surface. In 22 mo mice, while 2d labels were minimally affected by ROI, 16d labels consistently showed higher percentages in the medial ROI. Data are reported as percentages (labeled lacunae/all lacunae). Boxplots represent mean value (cross), interquartile range (box), minimum/maximum (whiskers), and symbols representing all data points. All p-values correspond with results of the omnibus ANOVA test.

We also divided each ROI into three sections with respect to the position between the endocortical and periosteal surface, since tissue strain increases with distance away from the centroid. Distance sections included 0-30%, 30-70%, and 70-100% (**Figure 7B**). There were not significant interactions between ROI and distance for either 5 mo or 22 mo mice. In 5 mo mice, both 2d-labeled and 16d-labeled lacunae percentages were influenced by the intracortical position but not by ROI. At this age, posthoc testing reveals that the lacunae closest to the periosteal surface (last 70-100% of cortical thickness) had a smaller percentage of 2d labels compared to the other two distances (**Figure 7C**). There were fewer 16d-labeled lacunae within the middle distance (30-70% of cortical thickness) compared to the other distances (**Figure 7D**). Differences in the percentage of labeled lacunae with intracortical position were more pronounced for 22 mo mice. At this age, lacunae within the middle intracortical distance had lower percentage of 2d labels compared to other the other distances (**Figure 7E**). For 16d labels, lacunae near the periosteal surface showed a higher 16d label percentage compared to the other distances (**Figure 7F**).

### 3.6 Osteocytes with active LCS turnover have larger lacunae

We conducted a quantitative analysis of backscattered SEM images at the anterior ROI of femoral cross-sections to assess whether lacunar geometry changes with aging and the recency of labeling. Since lacunar geometry differs between lamellar and non-lamellar compartments, and the relative size of these compartments changes in aging, we restricted our analyses to lamellar bone. Compared with 5 mo mice, 22 mo mice had decreased cortical lacunar porosity (–27%, p=0.037) but not lacunar number density (p=0.35) (**Figure S4**). Older mice also had smaller lacunae, as seen by smaller lacunar area (–22%, p=0.02), major axis (–12%, p=0.045), minor axis (–15%, p=0.05) (**Figure 8A, Figure S4**). No changes were seen in lacunar circularity with age. It is noted that previous work found that 2D SEM analysis is insufficient to detect expected increased sphericity in lacunae with aging.^62^

**Figure 8.**
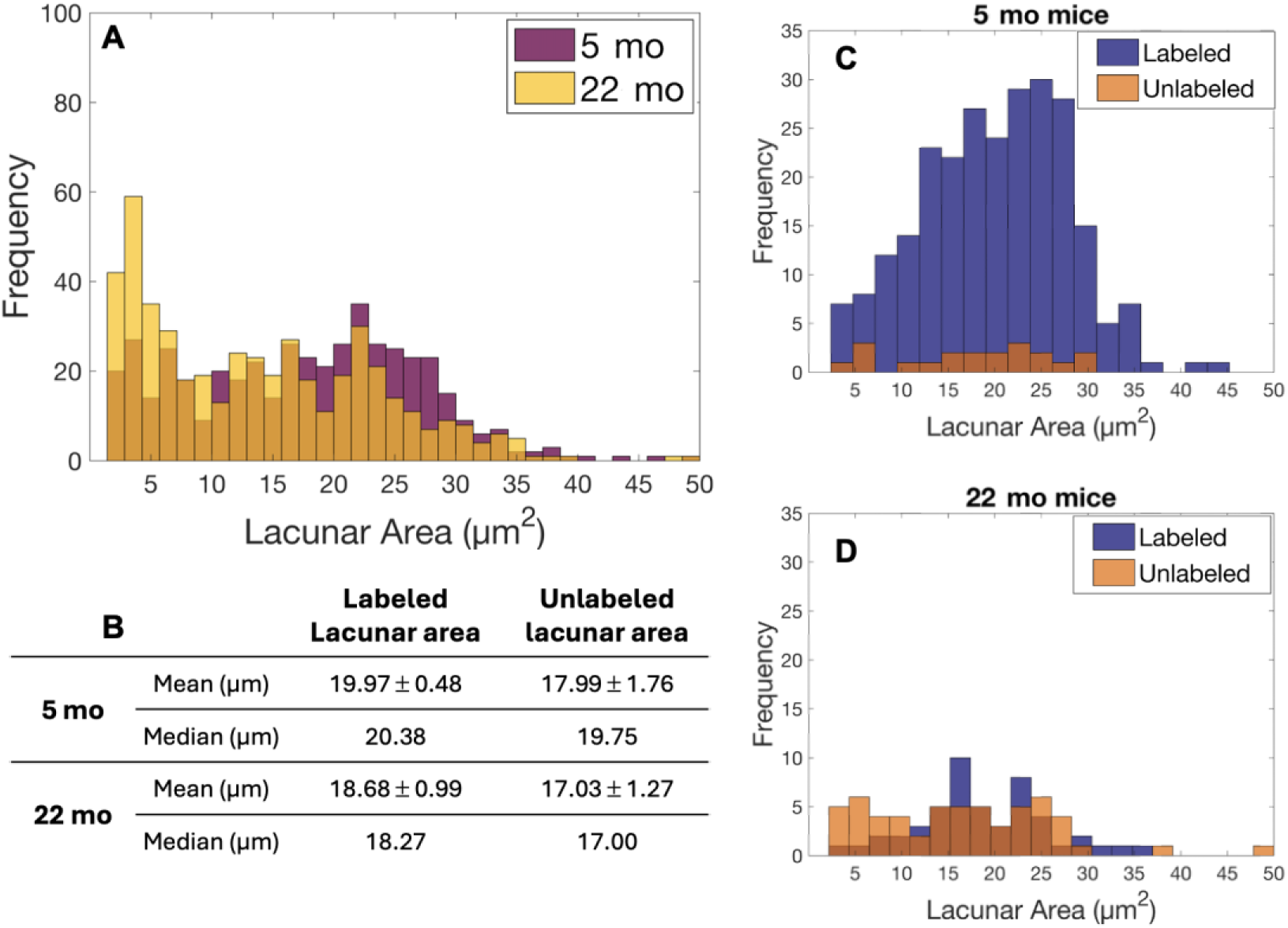
SEM analysis of lacunar geometry. A-B) labeled lacunae have larger areas compared to unlabeled lacunae in both ages and that C-D) the distribution of labeled lacunar sizes is approximately normal in both ages, whereas unlabeled lacunae show closer to a uniform distribution for both ages.

We also asked whether lacunar size differs between labeled and unlabeled lacunae by overlaying SEM and CLSM maps of the anterior region at the cortical midshaft femur. **Figure 8C** and **D** demonstrate that the distribution of labeled lacunar sizes is approximately normal for both 5 mo and 22 mo mice. By contrast, the sizes of unlabeled lacunae show closer to a uniform distribution for both ages. Our data suggest that recent LCS turnover increases lacunar area (**Figure 8 A & B**). Labeled lacunae had larger mean (+11% in 5 mo and +10% in 22 mo) and median areas compared with unlabeled lacunae.

## 4.0 Discussion

Osteocyte lacunar-canalicular system (LCS) turnover has been of high research interest as a possible contributor to age-related changes in bone fracture resistance^3,4,17,18,34,63,64^. Extensive evidence indicates that age reduces osteocyte viability and mechanosensitivity^7,8,14,25,65–69^ and truncates lacunar and canalicular dimensions and connectivity^20,21,62,70–73^. These changes imply that there is a decrease in LCS turnover in aging, which could have impacts to osteocyte mechanosensitivity and bone matrix quality^17,19,20,30,35,42,74^. However, currently, more questions than answers exist about where and how often osteocytes remove and replace their surrounding bone. Our study aimed to test the hypothesis that fewer osteocytes participate in LCS bone mineralization and resorption in aging C57BL/JN female mice (5 mo vs 22 mo). In this work, we find that aging reduces cortical and cancellous osteocyte participation in both perilacunar bone mineralization and resorption, in a manner that likely depends on tissue strain.

The osteocyte is known to remove and replace bone surrounding the LCS in response to perturbations in calcium homeostasis (e.g., lactation, PTH)^24,27,31,63^, but the participation of osteocytes in LCS turnover outside of calcium pressure has been uncertain^17^. We utilized serial fluorochrome labeling to estimate where, when, and how often osteocytes turn over their surrounding bone. Each mouse in this study was administered two fluorochrome labels at different times before euthanasia. We find that osteocyte bone turnover is highly prevalent in young adult mice, in both cortical and cancellous bone. Over 80% of osteocytes show fluorochrome labeled lacunae administrated 2 days before euthanasia. These numbers decrease to around 45% at 16 days before euthanasia, suggesting rapid bone turnover (**Figure 5**). Since label disappearance is an indirect indicator of bone resorption, we stained decalcified sections from the contralateral femurs for MMP14. In 5 mo mice, a comparable percentage of osteocytes are positive for bone resorption (i.e., MMP14-positive) as for bone mineralization (i.e., fluorochrome-labeled) (**Figure 4**). Together, these data suggest that osteocytes in the young adult murine skeleton engage in a pattern of frequent, near-daily bone mineralization along the LCS, coupled with frequent bone resorption events.

Aging has established deleterious impacts on the osteocyte. With increased age, osteocyte apoptosis and senescence both increase while autophagy decreases^8,14,69,75^. Aging reduces the size of lacunae and canaliculi, as well as the connectivity of this network^20,21,62,70–73^. These changes imply, but do not determine, that LCS turnover also changes with age. Here, we report that aging reduces LCS bone mineralization and, to a lesser extent, resorption. Compared with 5 mo mice, 22 mo mice have a 58% decrease in the percentage of 2d labeled lacunae and a 10% decrease in the percentage of MMP14+ lacunae in cortical bone (**Figure 4**). The rate of decrease in label percentage from 2d to 16d in cortical bone is similar across ages. These data suggest that while the number of osteocytes participating in LCS turnover decreases with aging, active osteocytes of different-aged cortical bone may have a characteristic time span of bone deposition before resorption events. These characteristics did not differ between lamellar and non-lamellar bone, demonstrating that the impact of aging on LCS bone turnover is not confounded by changes in cortical bone organization over the lifespan.

Osteocytes reside within cortical and cancellous bone but may have distinct roles within each of these compartments and different aging-induced impacts in their behaviors. There was a smaller age-related decline in the percentage of osteocytes engaged in LCS bone mineralization in cancellous versus cortical bone (–31% vs –58%, **Figure 4**). While the dynamics of bone turnover did not change largely with age in cortical bone (i.e., comparable rate of decrease in labeled lacunae from 2d to 16d between ages), age greatly increased the frequency of bone turnover in cancellous bone (**Figure 5**). These data suggest that more osteocytes remain active in cancellous bone and increase their rate of bone turnover compared with cortical bone. Cancellous bone is known to be more metabolically active than cortical bone^36–40^. It is possible that cancellous osteocytes have increased burden of participating in calcium homeostasis in the aging skeleton, but this hypothesis remains speculative at this time.

Our data suggest that the smaller lacunae reported in numerous imaging studies of the aging skeleton are associated with decreased osteocyte bone turnover. From coupling scanning electron microscopy measurements of lacunae with fluorochrome labeling, we find that osteocytes that are engaged in recent bone mineralization, regardless of the age group, reside within larger lacunae compared to osteocytes without labels (**Figure 8**). This result suggests that bone resorption events remove a considerable quantity of bone. It is yet to be fully determined which specific additional factors osteocytes employ to promote, or inhibit, local bone mineralization and/or formation. For example, it is well established that osteoblasts participate in local bone mineralization through shuttling hydroxyapatite precursors in vesicles to be released near forming surfaces^76^. Whether osteocytes engage in these active mineralization, or mineral inhabitation processes, should be investigated.

The influence of aging on osteocyte LCS turnover may be associated with tissue strain. In young adult mice, osteocyte participation in LCS turnover was abundant and did not differ between regions of the cortical femur with different tissue strain environments. By contrast, in early old age mice, osteocyte LCS bone mineralization varied between regions with different strain. Compared with osteocytes in anterior and posterior regions of cortical bone (i.e., higher tensile and compressive strains under loading, respectively^53–55^), more osteocytes in the medial region of aged bones (i.e., closer to neutral axis) were engaged in LCS bone mineralization and the LCS turnover rate was decreased (**Figure 7**). Similarly, in aged mice, the position of lacunae within the cortical thickness had a stronger relationship with LCS bone mineralization than for younger mice. For the 22 mo mice, the greater abundance of 16d labels in the bone closest to the periosteal edge may indicate that strain may influence bone resorption near osteocytes. While these data suggest a strain dependence on the LCS turnover behavior of osteocytes, whether there is a minimum strain to engage osteocyte LCS turnover, and whether this strain threshold changes in aging, would benefit from further investigation.

With aging, osteocytes become less mechanosensitive^9–16^. A persistent question is why aging has this effect on these long-lived cells. It has been recognized for many years that substantial strain amplification must occur in order for osteocytes to respond with anabolic signals to normal skeletal loads^43,77,78^. The changes in lacunar and canalicular shape with age may reduce tissue strain and fluid flow shear stress to contribute to these age-related changes in strain experienced by the cell, as shown by several finite element modeling studies ^45–48^. Our data add to this understanding by showing that changes in osteocyte lacunar size in age are approximately bimodal in distribution (i.e., only some aged osteocytes have much reduced lacunar size). Further, labeled osteocyte lacunae are larger than non-labeled lacunae at both ages.

These results suggest that contributions to osteocyte mechanosensation derived from geometric factors (i.e., shape of lacunae and canaliculi and the impact of these shape changes on fluid flow) is likely influenced by the activity of these cells in turning over their local bone. Additionally, our prior work demonstrated that labeled osteocyte lacunae are surrounded by more compliant (i.e., lower modulus) bone^34^. Thus, changes to LCS turnover in aging have multiple potential avenues of altering strain experienced by osteocytes. Our result that aging decreases the percentage of osteocytes engaged in LCS bone mineralization and resorption, but not the apparent rate of label disappearance (i.e., an estimate of resorption), may align with data from studies on the impact of aging on calcium signaling. In the cortical bone of 22 mo female C57BL/6JN mice, there are fewer osteocytes (∼ –60%) with active calcium signaling compared to younger mice, yet the remaining osteocytes respond to mechanical load with Ca^2+^ peaks of comparable intensities to those observed in young mice^9^. It is not yet understood whether populations of aged osteocytes with different LCS bone turnover characteristics vary in their mechanosensitivity. Together, these data suggest that major gaps still exist in our understanding about the strain experienced by the osteocyte and how these strains change in aging.

Our data also add to the emerging understanding of the osteocyte as a cell with the potential to directly modify bone matrix properties. As demonstrated in transgenic mouse studies, mice with decreased ability to engage LCS turnover have more fragile cortical bone^30,35,42^. While our earlier data demonstrates that LCS turnover increases perilacunar bone compliance in both young (5 mo) and aged (22 mo) mice^34^, the scale of osteocyte participation in LCS turnover was unknown. Here, we show that far less bone is turned over by osteocytes with increased age. These data add evidence to the potential beneficial effect of LCS turnover on bone matrix quality. However, several important questions remain that deserve increased attention. Specifically, how the osteocyte impacts local bone matrix is uncertain. Whether osteocytes specifically form, shape, or cross-link collagen are actively debated and would benefit from mechanistic investigations^17^.

There were several important limitations to this study. First, age-related changes in metabolic processes and LCS network architecture may impact fluorochrome dye uptake between cells in young and old bones and this limitation should be addressed in future studies. Additionally, our study focused solely on female mice, and the literature highlights important sex differences in osteocyte protein expression and potentially osteocyte metabolism and endocytosis of membrane proteins^41,79,80^. Moreover, extending the age range of the study would be beneficial to explore if LCS turnover changes during the developmental and advanced ages. In this study, it was not possible to investigate the age of individual osteocytes in older bones and discern whether the active osteocytes were young or old. Future investigations should also aim to assess whether changes in the shear stress, induced by interstitial fluid flow, play a significant role in the observed impacts of the strain environment, particularly the intracortical strain, on LCS turnover.

In summary, this study presents the first evidence that osteocyte participation in mineralizing their surroundings is highly abundant in both cortical and cancellous bone of young adult female C57BL/6JN mice. In aging, there are fewer osteocytes with active LCS turnover (both bone mineralization and resorption), yet turnover dynamics remains mostly similar in cortical bone of 5 mo and 22 mo mice, suggesting that active osteocytes engage in a characteristic LCS turnover response. Our results also demonstrate that the impacts of aging on LCS turnover are not the same at all locations throughout the femoral cortex and specifically differ with tissue strain. The large decline in LCS turnover in aging can have significant implications for bone quality, since osteocytes with active turnover have larger lacunae in both young and old bones as well as more compliant perilacunar tissue^34^. These results together signify a potential role for osteocyte bone turnover in the loss of bone fracture resistance and changes in mechanosensation in aging.

## Acknowledgments

This research was made possible by the Department of Mechanical & Industrial Engineering and the College of Engineering at the Montana State University. This work represents the views of the authors and not necessarily those of the sponsors. Research reported here is supported by NSF 2120239 and NIH R03AG068680. Imaging method development and data collection were made possible by the help of Dr. Heidi Smith from the Center for Biofilm Engineering imaging facility at Montana State University, supported by funding from the National Science Foundation MRI Program (2018562), the M. J. Murdock Charitable Trust (202016116), the US Department of Defense (77369LSRIP & W911NF1910288), and by the Montana Nanotechnology Facility (an NNCI member supported by NSF Grant ECCS-2025391). Steven Watson is thanked for his help with image processing and dissection. Kenna Brown and Connor Devine are thanked for their assistance with tissue harvest. Grace Roaming, Chloe Woodwall, and Shane Stauffer are thanked for their help with sample preparation. We thank Fatema Aljamal, Leah Davidson, Torie Prall, Megan Brenna, Kelly Silk, and Allison Stevens for their help with initial steps of method development for this project. We thank Montana State University’s Animal Resource Center staff and especially Tamara Marcotte for the help in providing excellent mouse care.

## Author Contributions

Ghazal Vahidi: Investigation; methodology; data curation; formal analysis; visualization; writing – original draft; writing – review and editing. Connor Boone: Investigation; methodology; writing – review and editing. Fawn Hoffman: Formal analysis; Investigation; methodology; writing – review and editing. Chelsea Heveran: Conceptualization; methodology; formal analysis; resources; writing – original draft; writing – review and editing

