## Supplementary information for "Aging decreases osteocyte lacunar-canalicular turnover in female C57BL/6 mice"

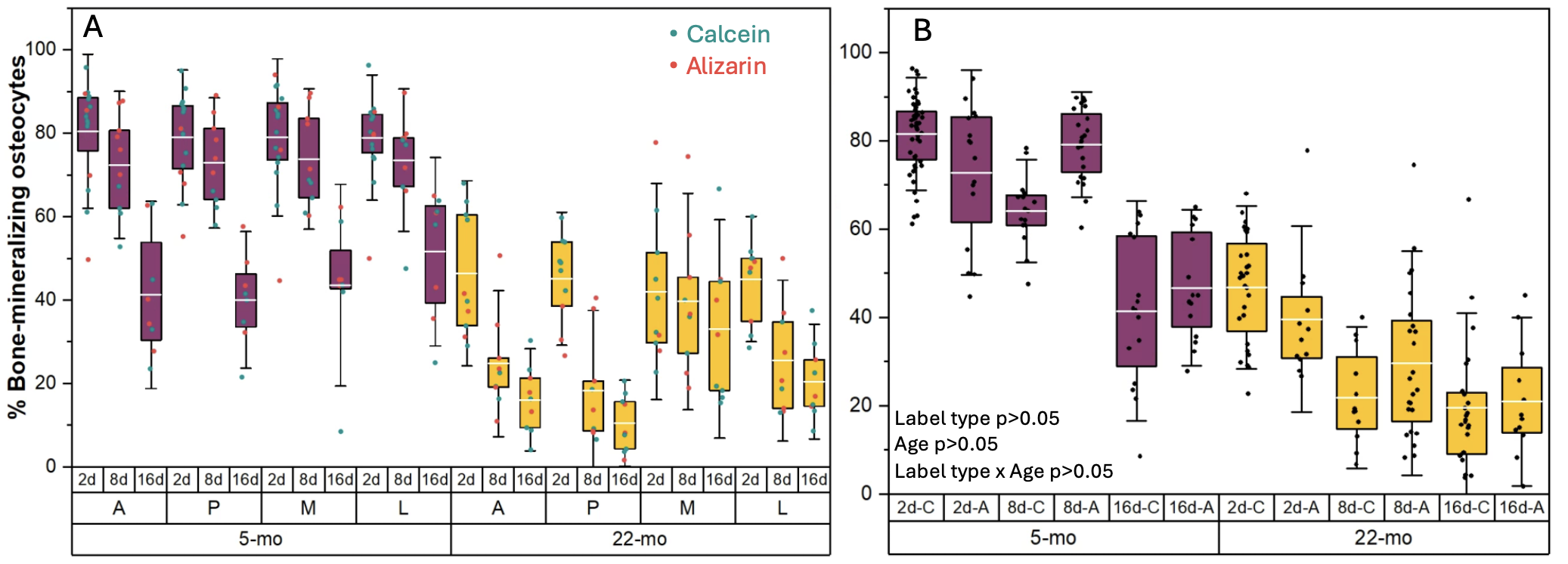

**Figure S1.** Fluorochrome label order does not influence the percentage of labeled lacunae. The order of calcein and alizarin labeling was alternated for a subset of mice to minimize the potential influence of label identity on measurements of labeled lacunae at each injection time point. For every combination of time points, some mice received the calcein injection first followed by the alizarin injection, while others received the labels in the reverse order. A) The percentage of bone-mineralizing lacunae (i.e., labeled lacunae) for the cortical bone of 5 mo and 22 mo mice at various ROIs and injection dates (2, 8, and 16d before euthanasia). The data points for calcein and alizarin are depicted in teal and red, respectively. B) Label order did not impact the percentage of bone mineralizing osteocytes for either age. Boxplots represent mean value (cross), interquartile range (box), minimum/maximum (whiskers), and symbols representing all data points. All p-values correspond with results of the omnibus ANOVA test.

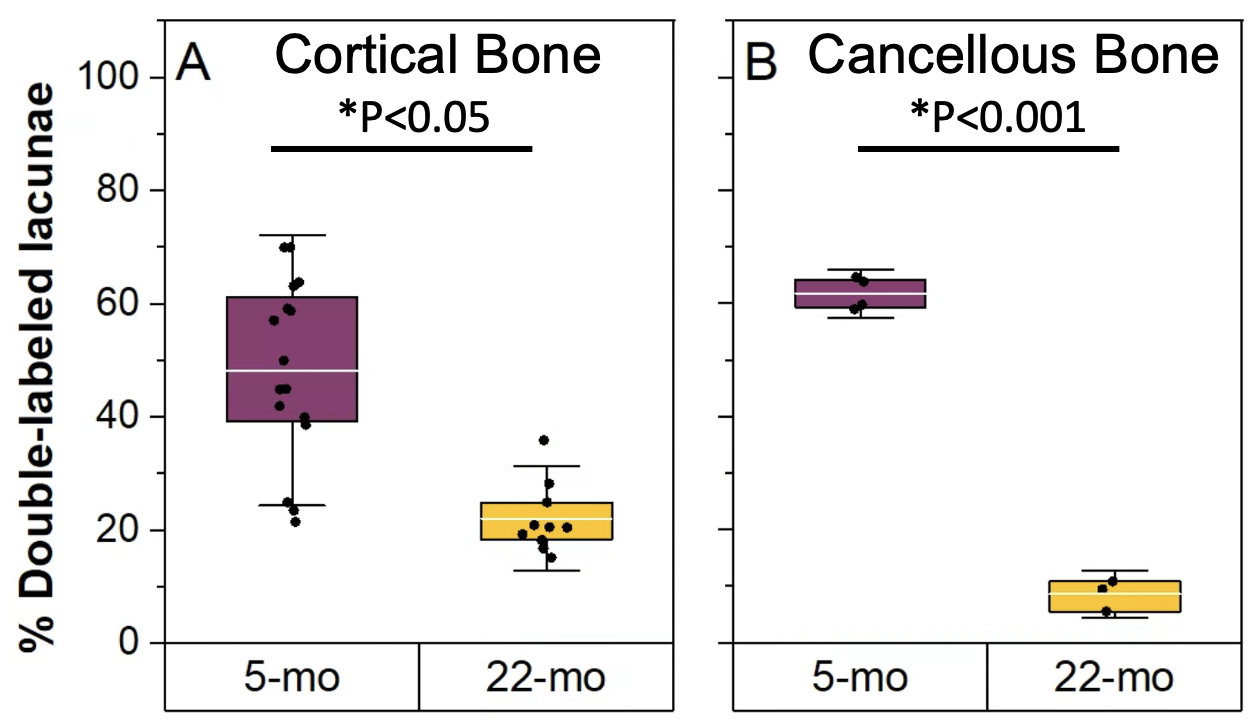

**Figure S2.** The effect of aging on the percentage of double-labeled lacunae in cortical and cancellous bone. **A *&* B)** In cortical and cancellous bone, double-labeled lacunae (i.e., had both 2d and 16d labels) were abundant in 5 mo mice. The percentage of double-labeled lacunae declined with age (22 mo vs 5 mo: cortical bone, -45% and p = 0.05; cancellous bone, -85% and p < 0.001). Boxplots represent mean value (cross), interquartile range (box), minimum/maximum (whiskers), and symbols representing all data points. All p-values correspond with results of the two-sided t-test. * represents significant age effect.

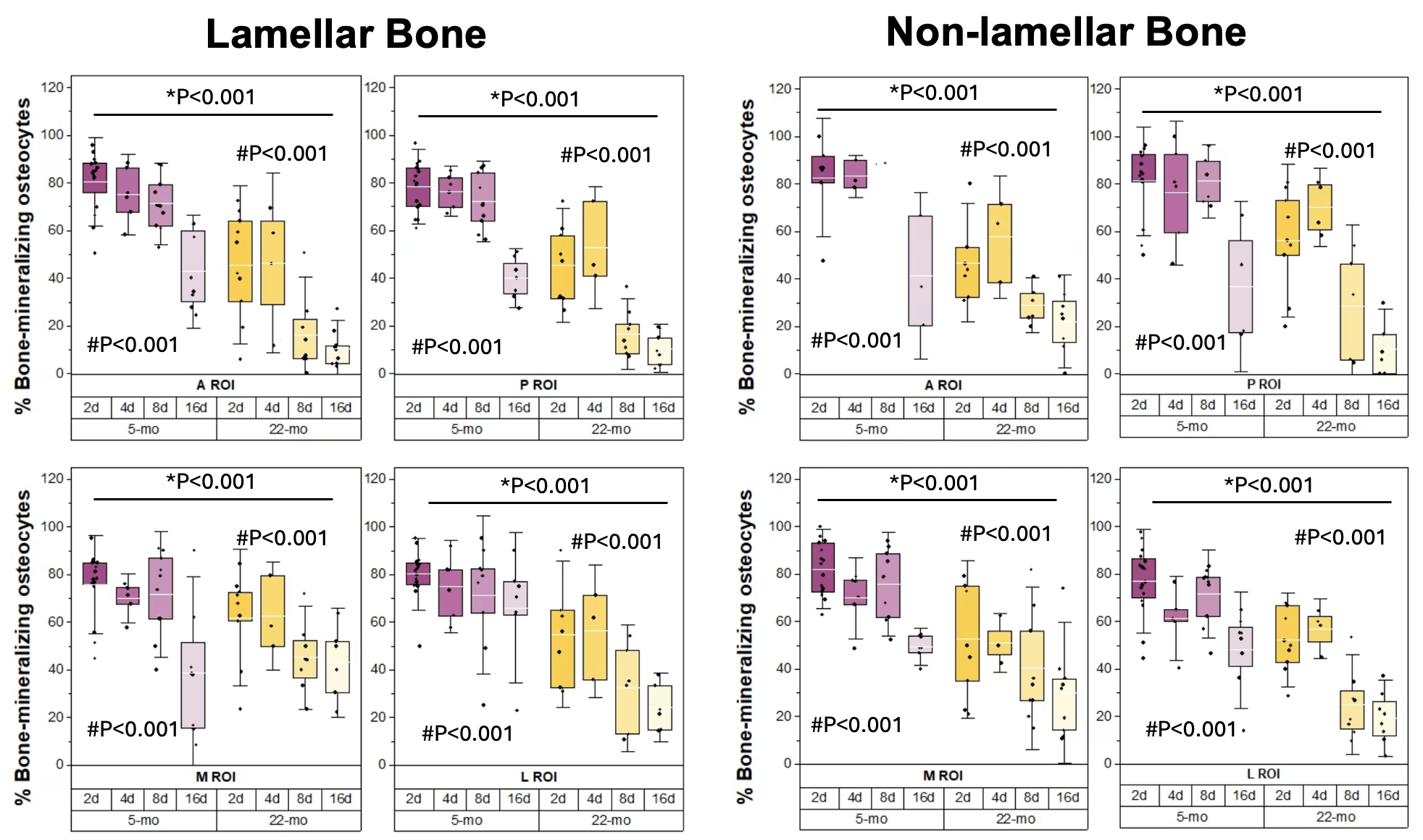

**Figure S3.** LCS bone mineralization in lamellar and non-lamellar bone. In all cortical ROI regions**,** aging reduced the percentage of bone-mineralizing osteocytes for both lamellar and non-lamellar compartments. For both type of tissues, 16d labeled lacunae were significantly less abundant compared to 2d labeled lacunae, regardless of the age group. All data are reported as percentages (labeled lacunae/all lacunae). Boxplots represent mean value (cross), interquartile range (box), minimum/maximum (whiskers), and symbols representing all data points. All p-values correspond with results of the omnibus ANOVA test. .* indicates a significant effect of age. # indicates a significant effect of injection date.

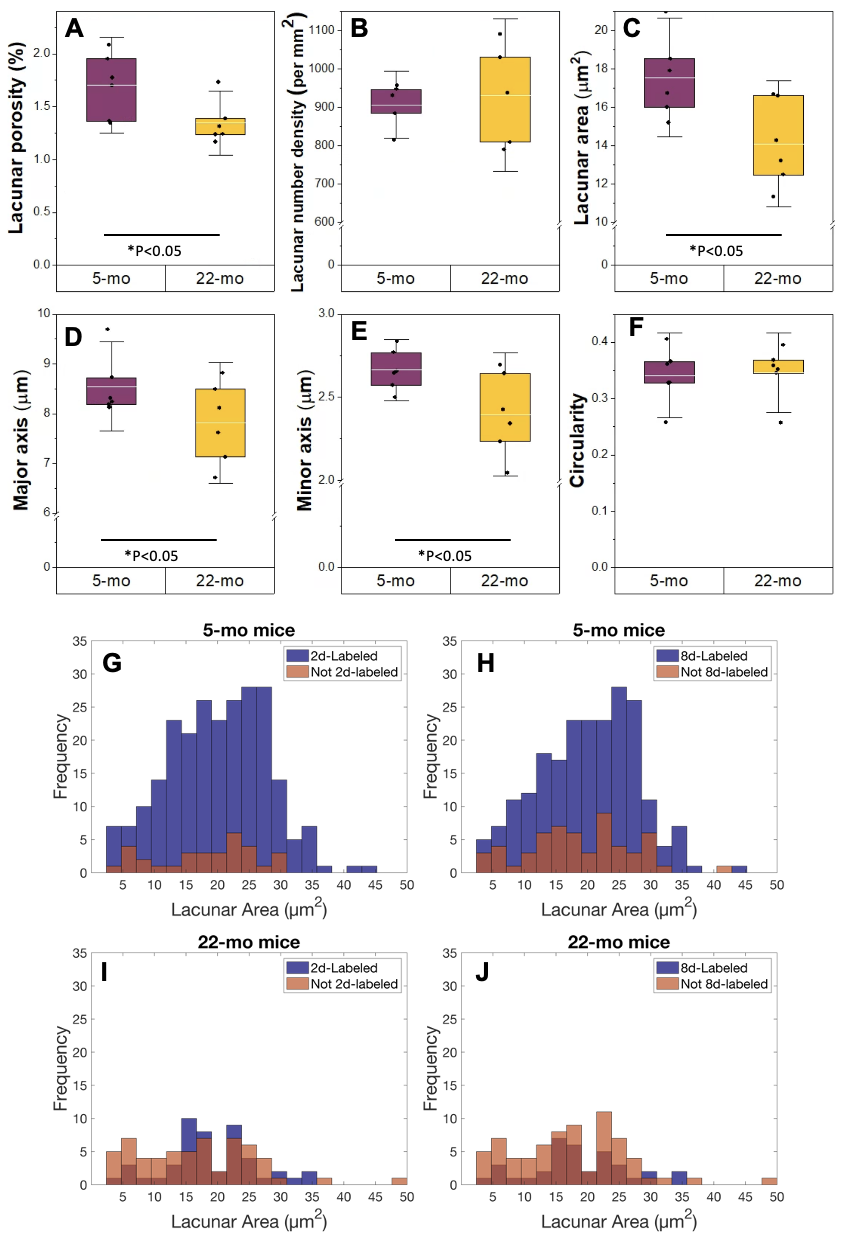

**Figure S4.** Scanning electron microscopy analysis of lacunar geometry. Aging A) increased lacunar porosity B) but did not change lacunar density. C) Lacunar area, B) lacunar major axis, and C) lacunar minor axis were smaller in cortical bone of 22 mo mice compared to 5 mo mice. D) Aging didn’t change lacunar circularity. We then analyzed lacunar size in labeled (i.e., 2d labeled and/or 8d labeled) and unlabeled (i.e., no 2d or 8d labels) lacunae by overlaying SEM and CLSM maps of the anterior cortical femur. G-J) In both ages, the distribution of lacunar sizes for both 2d and 8d labels was close to a normal distribution, whereas the distribution of lacunar sizes for unlabeled lacunae, especially in 5 mo bones, was closer to a uniform distribution. Boxplots represent mean value (cross), interquartile range (box), minimum/maximum (whiskers), and symbols representing all data points. All p-values correspond with results of the omnibus ANOVA test. .* represents significant age effect.
